## Supplemental Figures and legends for "Histone variants shape chromatin states in Arabidopsis"

**Figure 1 figure supplement 1.** Biochemical analysis of the association between histone variants and histone marks. (**A**) Analysis of DNA in input samples used for H3.1 and H3.3 mononucleosome immunoprecipitation after MNase digestion. (**B**) Silver-stained 8-20% gradient gel of immunoprecipitated H3.1 and H3.3 mononucleosomes. Arrows indicate transgenic (T) and endogenous (E) H3 proteins. Molecular weight markers are indicated on the right. (**C**) Analysis of DNA extracted from immunoprecipitated H3.1 and H3.3 mononucleosomes. (**D**) Spectral counts of H3.1- and H3.3-specific peptides (as indicated in panel **E**) in mononucleosomes immunoprecipitated with H2A variant-specific antibodies. (**E**) H3.1- and H3.3-specific peptides used for the analysis of H3K27, H3K36, and H3K37 modifications. (**F**) Number of MS measured spectra that covered the indicated residues in H3.1 and H3.3 mononucleosomes. (**G**) Number of measured MS spectra containing the indicated modifications in H3.1 and H3.3 mononucleosomes. (**H**) (top) Relative methylation levels of H3K9me1 and H3K9me2 on H3.1 and H3.3 revealed from transgenic copies of the respective H3 variants. (bottom) Sequences of peptides used to evaluate H3K9me1 and H3K9me2 by MS. (**I**) Sequences of peptides used to evaluate H3 acetylation by MS. (**J**) Relative acetylation levels of the indicated H3 residues in immunoprecipitated H3.1 and H3.3 mononucleosomes without differentiating H3.1 and H3.3 variants. (**K**) Relative acetylation levels of the indicated H3 residues on H3.1 and H3.3 revealed from transgenic copies of the respective H3 variants.

**Figure 2 figure supplement 1.** Validation of H2A.2 and H2A.Z.11 polyclonal antibodies. (**A**) Sequence alignments of three H2A.Z and four H2A histone variants. Positions that differ between variants are highlighted in blue. Underlined are sequences of peptides used for immunization. (**B**) Antibodies against H2A.2 were tested with four H2A variants expressed in bacteria. Relevant part of Coomassie stained gel with overexpressed H2As is shown on the top. As shown on the bottom panel antibody raised against H2A.2 peptide recognized only H2A.2. Antibodies against H2A.Z.11 were tested on nuclear extracts from WT, *h2a.z.9* knock-down/*h2a.z.11* knock out (*h2a.z.9 KD* *h2a.z.11* *KO*; this line still expresses H2A.Z.8), and *pie1-3* (mutant in large subunit of the SWR1 complex responsible for deposition of H2A.Z) lines. The H2A.Z.11 antibodies did not recognize any protein in the *h2a.z.9 KD* *h2a.z.11* KO line indicating its specificity. In *pie1-3* line, both H2A.Z.9 and H2A.Z.11 were detected but at reduced levels as compared to WT which is in line with the inefficient deposition of H2A.Z in the absence of SWR1. The H3 antibody served as the loading control. (**C**) Immunostaining of WT nuclei with indicated antibodies.

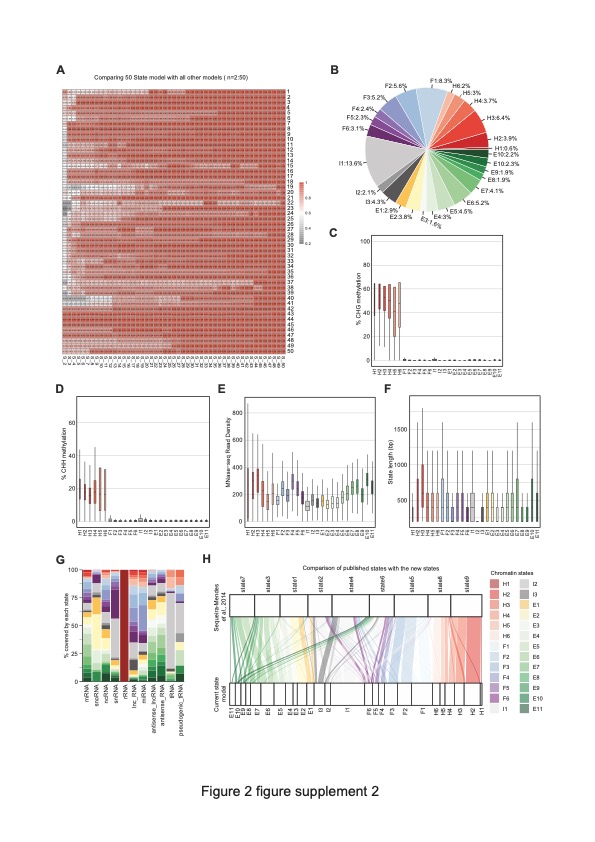

**Figure 2 figure supplement** **2.** Properties of chromatin states in *Arabidopsis thaliana.* (**A**) Heatmap showing the correlation (0 to 1) between emission parameters of each model. The Y axis represents the distinct states in the reference state model (n=50). The X-axis represents models including 2 to 49 states compared to the 50-state model. The correlation plotted here is the maximum correlation between each state of the reference model and any state of the other models being compared. (**B**) Pie chart showing the proportion of the genome covered by each state. (**C, D**) Box plots showing levels of CHG (**C**) and CHH (**D**) methylation across states. (**E**) Box plot showing nucleosome occupancy (MNase-seq read density) across states. (**F**) Box plots showing the length (bp) of chromatin states. (**G**) Stacked bar plot showing the overlap between non-protein-coding gene features and chromatin states. (**H**) Flow diagram showing the overlap (in bp) between published states (Sequeira-Mendes et al., 2014) and the current states. The color code for the flow diagram represents the chromatin states in the current model.

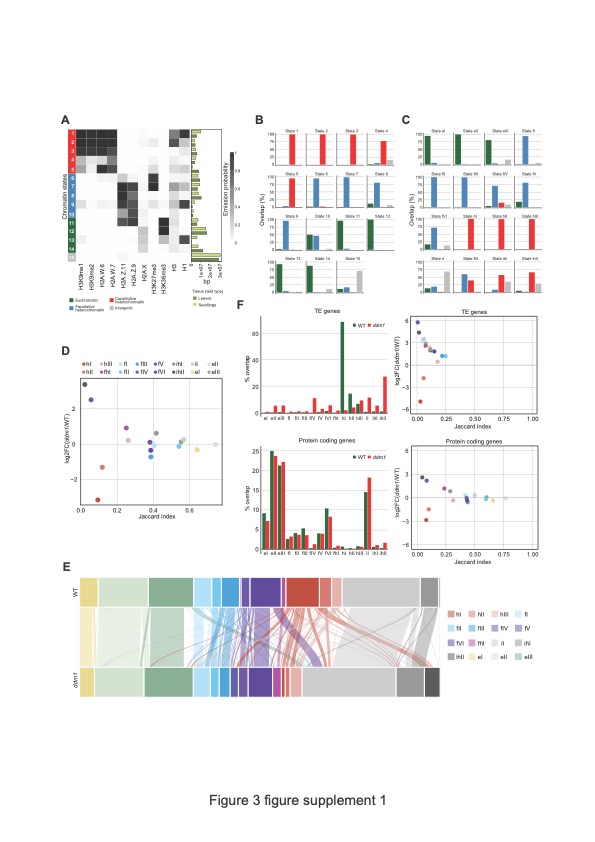
 **Figure 3 figure supplement 1.** DDM1 loss of function disrupts chromatin states in *Arabidopsis thaliana.* (**A**) Heatmap showing the emission probability of histone mark/variant across the 15 chromatin states of the concatenated model computed with chromatin profiles from seedlings and leaves. The bar plot on the right represents the proportion of the genome covered by each state in seedlings (light green) and in leaves (dark green). (**B**) Bar plots of the genomic overlap between the seedling and leaf concatenated model states and states from the four chromatin types in the 26-state model. 14 states show overlaps with predominantly one chromatin type, whereas only state 10 overlaps with two types in similar proportions. (**C**) Bar plots of the genomic overlap between the wild type and *ddm1* concatenated model states and states from the four chromatin types in the 26-state model. 13 states overlap with predominantly one chromatin type, whereas 3 states overlap with two types. In **B** and **C**, color codes for four major groups of chromatin states are as in panel **A**. (**D**) Scatter plot of the Jaccard index *vs*. the log2 fold change in proportion of genome for the 16 states from the wild type and *ddm1* concatenated model. For each state the Jaccard index was calculated for the genomic regions covered in wild type and in *ddm1*. The log2 fold change was calculated using the total number of base pairs covered by each state in *ddm1* mutant compared to in the wild type (see also methods). (**E**) Alluvial plot showing the chromatin state changes between the wild type (top row) and *ddm1* (bottom row). The plot was generated using the R package ggalluviel v0.12.5 (<http://corybrunson.github.io/ggalluvial/>). (**F**) Scatter plots of the Jaccard index *vs.* the log2 fold change in proportion of genome for the 16 states from the wild type and *ddm1* concatenated model (see **Figure 3 figure supplement 1D** and Methods) for TE genes and protein coding genes separately (right panels). In addition, the percentages of the combined length of the TE/protein coding genes covered by each state are shown as bar plots on the left. Color codes for chromatin states are as in panel **D**.

**Figure 3 figure supplement 2.** Interaction of DDM1 and DDM1 deletion mutants with histone variants. (**A-C**) Identification of H2A.W and H2A.Z binding sites in DDM1. His_6_- (**A**) or GST- (**B** and **C**) tagged DDM1 fragments, as indicated with amino acids numbers, were incubated with indicated H2A-H2B dimers and after washing samples were analyzed on 15% SDS-PAGE and stained with Coomassie.

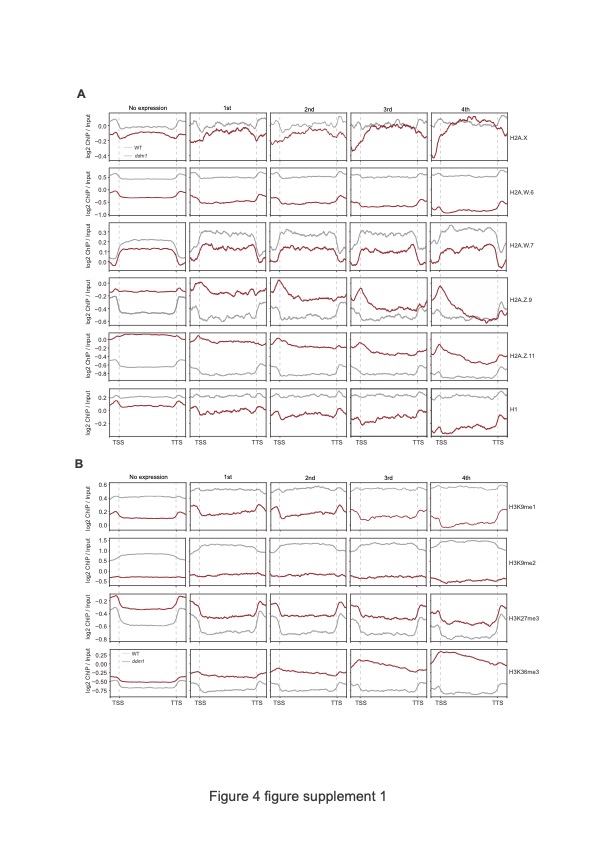

**Figure 4 figure supplement** **1.** Analyses of the parameters that could correlate the chromatin states with TE expression in *ddm1.* Enrichment profiles of histone variants (**A**) and H3 modifications (**B**) plotted over groups of TE genes based on their expression in *ddm1* as defined in the legend for **Figure 4A** and Methods.

**Figure 4 figure supplement 2.** Analyses of the parameters that could correlate the chromatin states with TE expression in *ddm1.* (**A**) Box plot showing the expression of the TE genes grouped by the state overlapping the TSS in *ddm1*. See also **Figure 4C**. (**B**) Box plot showing the TE gene length distributions across the 5 different TE gene expression groups. (**C**) Box plots showing state statistics for TE genes per expression group in *ddm1* and wild type (WT). “# state types” (top) is the number of different kinds of states present over the TE genes. “# state changes” (middle) is the number of times there is a transition from one state to another and the bottom shows the length of the states overlapping the TE genes. No trend can be seen across expression groups, whereas all TE genes tend to have a larger diversity of states, more frequent transitions between states and hence shorter stretches of each state.
